# Novel inhibitors of acute, axonal DLK palmitoylation are neuroprotective and avoid the deleterious side effects of cell-wide DLK inhibition

**DOI:** 10.1101/2024.04.19.590310

**Authors:** Xiaotian Zhang, Heykyeong Jeong, Jingwen Niu, Sabrina M. Holland, Brittany N. Rotanz, John Gordon, Margret B. Einarson, Wayne E. Childers, Gareth M. Thomas

## Abstract

Dual leucine-zipper kinase (DLK) drives acute and chronic forms of neurodegeneration, suggesting that inhibiting DLK signaling could ameliorate diverse neuropathological conditions. However, direct inhibition of DLK’s kinase domain in human patients and conditional knockout of DLK in mice both cause unintended side effects, including elevated plasma neurofilament levels, indicative of neuronal cytoskeletal disruption. Indeed, we found that a DLK kinase domain inhibitor acutely disrupted the axonal cytoskeleton and caused vesicle aggregation in cultured dorsal root ganglion (DRG) neurons, further cautioning against this therapeutic strategy. In seeking a more precise intervention, we found that retrograde (axon-to-soma) pro-degenerative signaling requires acute, axonal palmitoylation of DLK and hypothesized that modulating this post-translational modification might be more specifically neuroprotective than cell-wide DLK inhibition. To address this possibility, we screened >28,000 compounds using a high-content imaging assay that quantitatively evaluates DLK’s palmitoylation-dependent subcellular localization. Of the 33 hits that significantly altered DLK localization in non-neuronal cells, several reduced DLK retrograde signaling and protected cultured DRG neurons from DLK-dependent neurodegeneration. Mechanistically, the two most neuroprotective compounds selectively prevent stimulus-dependent palmitoylation of axonal pools of DLK, a process crucial for DLK’s recruitment to axonal vesicles. In contrast, these compounds minimally impact DLK localization and signaling in healthy neurons and avoid the cytoskeletal disruption associated with direct DLK inhibition. Importantly, our hit compounds also reduce pro-degenerative retrograde signaling in vivo, suggesting that modulating DLK’s palmitoylation-dependent localization could be a novel neuroprotective strategy.

## Introduction

Dual-leucine zipper kinase (DLK) is an upstream activator (a ‘MAP3K’) of mitogen-activated protein kinase (MAPK) pathways that is highly expressed in neurons. DLK functions as an evolutionarily conserved ‘stress sensor’ (1–4) and one major role of DLK is to convey retrograde signals from sites of axonal damage or disruption back to neuronal nuclei (5–10). Genetic or pharmacological block of DLK inhibits such retrograde signaling, and is neuroprotective, after trophic factor deprivation (TD) in developing neurons, as well as in models of acute injury and of neurodegenerative disease in the mature nervous system (5, 6, 10–12). Importantly, DLK was also initially reported to selectively mediate pro-degenerative, but not basal, physiological signaling by downstream MAPKs (5). These findings suggested that inhibiting DLK could be a promising therapeutic strategy to prevent diverse forms of neurodegeneration and spurred development of inhibitors of DLK’s kinase activity (13–15). Based on these and other encouraging preclinical findings, one such DLK inhibitor was moved forward to a Phase I clinical trial (13, 16). However, numerous patients in this trial developed symptoms indicative of sensory neuropathy which, along with other adverse events, led a significant proportion of those enrolled to reduce or cease dosage (16). Further analysis revealed that DLK inhibitor treatment elevated levels of neurofilament in patients’ plasma, suggestive of axonal cytoskeletal disruption (16). This effect is likely due to on-target toxicity as plasma neurofilament levels were also elevated when DLK was conditionally and globally knocked out in adult mice (16). Although it was unclear if this elevation was due to a neuron-intrinsic role of DLK, these findings suggested that broadly inhibiting DLK’s kinase activity causes deleterious side effects, cautioning against such a therapeutic approach.

We report here that the disruption of the axonal cytoskeleton caused by DLK inhibition is indeed likely to be neuron-intrinsic, as evidenced by similar outcomes after acute treatment of cultured DRG neurons with a DLK inhibitor. We therefore considered the possibility that more refined strategies to inhibit specific pools of DLK, particularly those involved in pro-degenerative retrograde signaling, might afford neuroprotection without cytoskeletal disruption. In particular, our prior work revealed that DLK is covalently modified with the lipid palmitate (8). This modification, palmitoylation, targets DLK to lipid membranes and in neurons is critical for DLK to ‘hitchhike’ on axonal trafficking vesicles (8). Palmitoylation is also critical for DLK’s function; by genetically inhibiting DLK palmitoylation through the use of shRNA knockdown and rescue to replace endogenous DLK with a palmitoyl-site mutant, we found that this approach blocks DLK-dependent retrograde signaling and exhibits neuroprotective effects, both in cultured neurons and in vivo.(8, 9, 17). Importantly, the protective effects of DLK palmitoyl-site mutation are as robust as complete loss of DLK (9, 17).

Genetically preventing DLK palmitoylation (e.g. by mutating DLK’s palmitoylation site at the genomic level, perhaps using Clustered Regularly Interspaced Short Palindromic Repeats (CRISPR)-based methods) is unlikely to be feasible therapeutically. However, the above findings suggested that pharmacologic block of palmitoyl-DLK-dependent signaling might serve as an alternate neuroprotective strategy. Building on this notion, we developed a high content imaging screen to identify compounds that alter the palmitoylation-dependent localization of transfected, GFP-tagged DLK (DLK-GFP) in non-neuronal human embryonic kidney (HEK) 293T cells (18). Wild type (wt) DLK-GFP normally localizes to Golgi membranes and small vesicle-like puncta in these cells, but pharmacologic or genetic block of DLK palmitoylation results in a diffuse DLK-GFP signal (8, 18). DLK-GFP’s punctate localization can thus be used as a proxy for DLK palmitoylation that is compatible with a High Content Screening (HCS)-based approach (18). A pilot screen using this assay identified a compound, ketoconazole, which reduced DLK-GFP’s punctate localization in HEK293T cells and also partially reduced acute DLK-dependent signaling in neurons shortly after TD (18). However, we reasoned that this partial, acute inhibition might not be sufficient to protect DRG neurons against degeneration induced by prolonged TD. We therefore sought to expand our screen using larger libraries, with the objective of identifying more potent compounds capable of preventing not just DLK-dependent signaling but also DLK-dependent neurodegeneration itself.

Here, we report the results of this screen, which identified 33 compounds (out of a primary screen of >28,000 compounds) that inhibit pro-degenerative DLK-dependent signaling in neurons. We further show that a subset of these compounds significantly protects DRG neurons against degeneration induced by prolonged TD. Mechanistically, we report that TD induces an acute palmitoylation-dependent change in the axonal localization of DLK that is prevented by our most neuroprotective hits, providing a plausible explanation for their neuroprotective activity. Crucially, our top hits do not phenocopy the disruption of the axonal cytoskeleton seen after DLK kinase inhibition. Moreover, intravitreal injection of our top hits significantly reduces DLK-dependent pro-degenerative retrograde signaling in the retina. These new classes of inhibitors thus reduce DLK-dependent pro-degenerative retrograde signaling without causing side effects associated with global DLK inhibition. This discovery reveals a novel therapeutic strategy aimed at mitigating neurodegeneration.

## Results

### A neuron-intrinsic role of DLK in governing cytoskeletal integrity in healthy axons

Treatment of human patients with the DLK kinase domain inhibitor GDC-0134, and cKO of DLK (gene name *Map3k12*) in mice, both cause elevated plasma neurofilament levels, a biomarker of cytoskeletal disruption and neurodegeneration (19). This finding suggests that DLK activity is required for normal structure and function of the axonal cytoskeleton. The association between GDC-0134 treatment and sensory neuropathy (16), a condition often linked to pharmacological or genetic disruption of the cytoskeleton of dorsal root ganglion (DRG) sensory neurons (20–23), further support this conclusion.

These phenotypes could be due to roles of DLK in a different subcellular compartment, or even in a different cell type, but we hypothesized that DLK might be intrinsically required for axonal cytoskeletal integrity in DRG neurons. To test this possibility, we assessed the effect of acute DLK inhibition on the distribution of Neurofilament heavy chain (NF-200) and neuron-specific βIII tubulin (Tuj1). In axons of healthy DRG neurons treated with vehicle, NF-200 and βIII tubulin were smoothly and uniformly distributed along axons. However, in cultures treated with the widely used DLK inhibitor GNE-3511 (structurally similar to GDC-0134 (12, 16)), NF-200 and Tuj1 signals were no longer smoothly distributed along axons but instead partly accumulated in axonal distortions (Fig 1A). Moreover, the vesicle marker VAMP2 also accumulated in these structures (Fig 1B), suggesting that the cytoskeletal disruption caused by DLK inhibition in turn, or in addition, disrupted axonal transport. DLK itself was also enriched in these accumulations (Fig 1A), suggesting that blockade of endogenous DLK activity at these sites is likely responsible for the observed distortions and accumulations. Quantified data for distribution of NF-200, Tuj1, DLK and VAMP2 are shown in Fig1C-F, respectively, with examples of image processing and analysis in Fig S1. Collectively, these findings uncover a previously unrecognized role for DLK in maintaining the integrity of axonal neurofilaments, microtubules, and/or vesicle-based transport in healthy axons.

**Figure 1.**
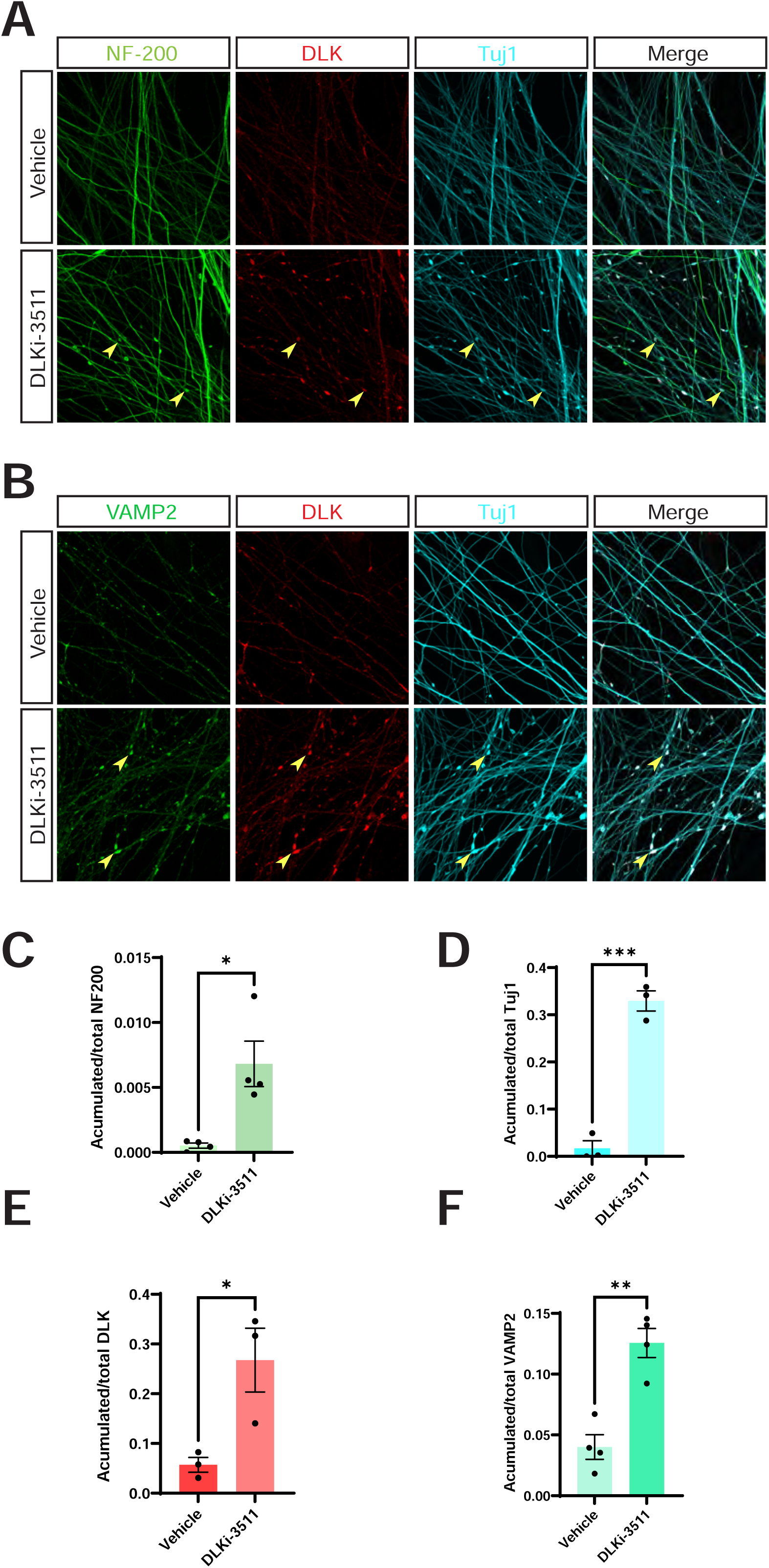
DLK kinase domain inhibition disrupts axonal integrity in DRG neurons. ***A:*** Images of cultured DRG neurons that had been treated with DMSO vehicle or 500nM DLK inhibitor GNE-3511 (DLKi-3511) for 4 hours and fixed and stained with the indicated antibodies. ***B:*** As *A*, except that neurons were fixed to detect vesicle marker VAMP2 rather than NF-200. ***C-F:*** Quantified data from *A* and *B* confirm that DLK inhibition disrupts NF-200 (neurofilament heavy chain; ***1C***) and Tuj1 (neuron-specific tubulin; ***1D***) distribution and leads to accumulations of DLK (***1E***) and VAMP2 (***1F***) in axons, suggestive of cytoskeletal dysregulation and/or impaired vesicle-based transport. Unpaired t tests reveal significant effects of DLKi-3511 vs. Vehicle treated in DLK signal (p =0.0330), Tuj1 signal (p =0.0003), NF-200 signal (p =0.0117), and VAMP2 signal (p =0.0016). In this and all subsequent panels, error bars represent the standard error of the mean (SEM).

### Palmitoylation-dependent recruitment of DLK to axonal vesicles is implicated in axonal retrograde signaling

These findings increase the likelihood that the peripheral neuropathy symptoms caused by DLK inhibition are due to dysregulation of cytoskeletal regulation and/or axonal transport in sensory neurons themselves. We therefore considered ways to specifically prevent pro-degenerative DLK signaling without globally inhibiting DLK. We previously reported that DLK must be palmitoylated to support axonal retrograde signaling (8, 9, 17), and a subsequent study reported that TD, which triggers pro-degenerative DLK signaling in developing DRG neurons (5), acutely increases DLK palmitoylation, and also induces DLK recruitment to axonal vesicles (24). However, the functional role of this acute, TD-induced change in DLK localization was not addressed. It was also unclear whether this change of DLK localization requires, or simply correlates with, increased DLK palmitoylation. As a first step to address these questions, we infected DRG neurons to express wild type, myc-tagged DLK (wtDLK-myc), subjected the neurons to TD and then immunostained to detect myc signal. TD increased wtDLK-myc recruitment to axonal puncta (Fig 2A, B), which based on prior work are likely vesicles (8, 24). We also investigated whether DLK recruitment to axonal vesicles involves acute palmitoylation of DLK itself. To address this question, we used ‘spot’ cultures, in which DRG axons extend from centrally plated cell bodies, thus allowing specific harvesting of axonal material. We subjected spot culture axonal fractions to a biochemical palmitoylation assay, acyl-biotin exchange (ABE) in which thioester-linked acyl groups (most commonly palmitate) are exchanged for biotin, allowing capture of the resultant biotinyl-proteins on avidin beads. Western blots of ABE fractions from DRG axons confirmed that TD acutely increases DLK palmitoylation (purity of axonal preparations for these experiments confirmed in Fig. S2). Importantly, a broad spectrum inhibitor of cellular palmitoylation, 2-bromopalmitate (2BP; (25)) prevented both the TD-induced increase in DLK palmitoylation and DLK recruitment to vesicles, increasing the likelihood that the former process drives the latter (Fig 2A, B and 2C, D). These findings confirm that TD triggers acute, axonal palmitoylation of DLK, and suggest that this acute palmitoylation is required for DLK to associate with axonal vesicles.

**Figure 2:**
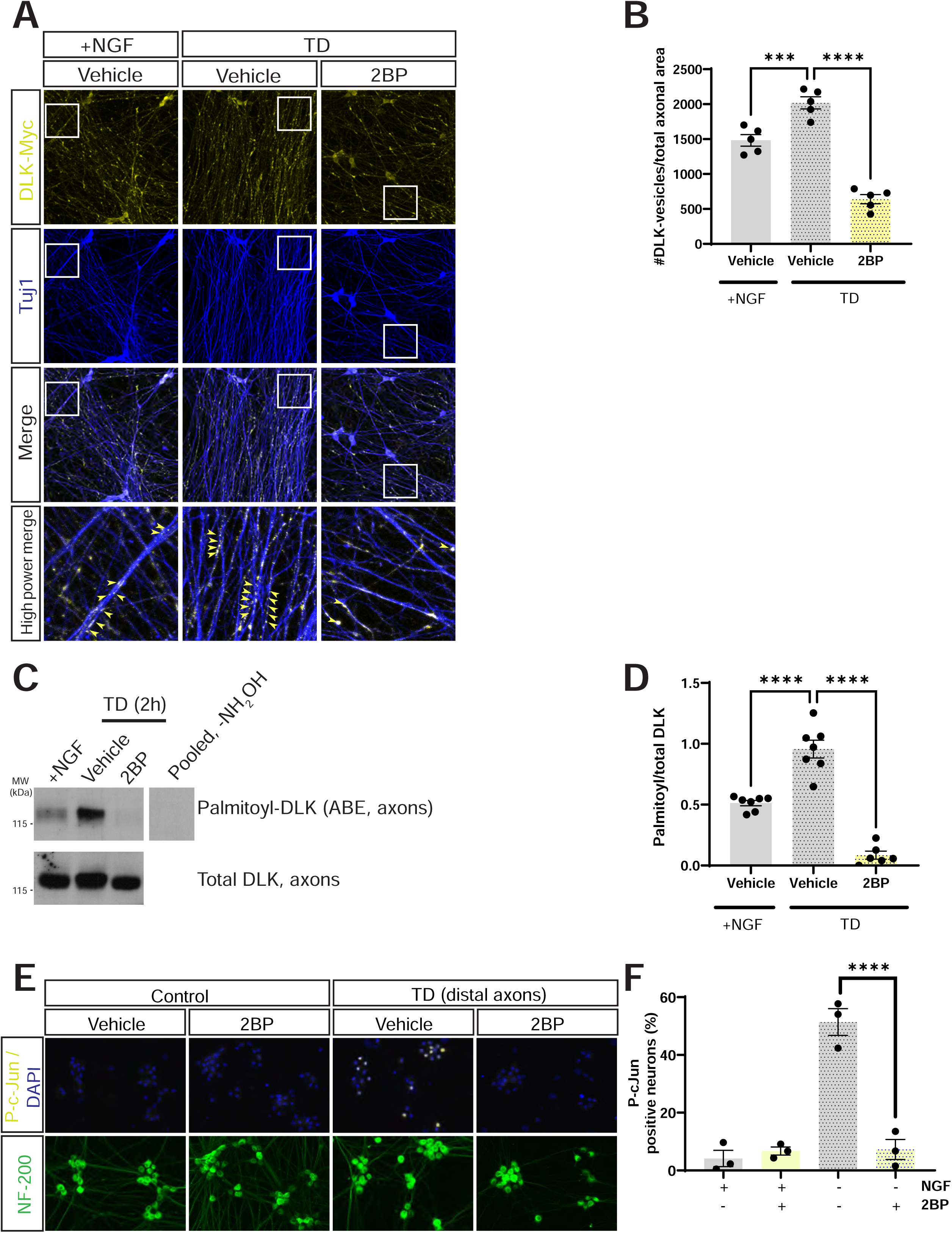
Trophic deprivation-induced recruitment of DLK to axonal vesicles and DLK-dependent retrograde signaling are both palmitoylation-dependent. *A:* Cultured DRG neurons were lentivirally infected to express myc-tagged wild type DLK (wtDLK-myc) and left untreated or were subjected to trophic factor deprivation (TD) in the presence of the indicated compounds or DMSO vehicle, prior to fixation and immunostaining with the indicated antibodies. The bottom row of images shows a magnified view of the boxed region in upper panels. *B:* Quantified data from *A* confirm that the TD-induced increase in axonal DLK puncta (presumptive vesicles (24)) is prevented by 2BP i.e. is palmitoylation-dependent. ***; p =0.0008; ****; p <0.0001, ANOVA, Dunnett’s post hoc test. *C:* Western blots of ABE (palmitoyl-) fractions of DRG axonal lysates that had been treated as indicated prior to lysis. The righthand lane is a side-by-side exposure from a parallel control sample omitting the key ABE reagent NH_2_OH, run on the same gel, with intervening spacer lanes cropped. *D:* Quantified data from *C* confirm that TD increases axonal DLK palmitoylation, which is prevented by 2BP. ****; p <0.0001, ANOVA, Dunnett’s post hoc test. *E:* Images of cell body chambers of DRG microfluidic cultures, fixed and stained with the indicated antibodies after selective treatment of distal axons as indicated. *F:* Quantified data from *E* confirm that TD-induced retrograde signaling requires acute palmitoylation in distal axons. ****; p<0.0001, ANOVA, Dunnett’s post hoc test.

To assess the functional importance of axonal palmitoylation for DLK-dependent retrograde signaling, we plated DRG neurons in microfluidic chambers and selectively withdrew trophic support from distal axonal chambers that had been treated with either 2BP or vehicle. Importantly, selective inhibition of axonal palmitoylation with 2BP prevented TD-induced c-Jun phosphorylation in DRG cell bodies (Fig 2E, F). This finding is consistent with a model in which acute, axonal palmitoylation of DLK is critical for TD-induced, pro-degenerative retrograde signaling. Importantly, these results also suggested that selectively preventing acute, palmitoylation-dependent recruitment of DLK to vesicles could be a novel neuroprotective strategy that might circumvent the side effects associated with inhibiting all cellular pools of DLK.

### High content screening to identify modulators of DLK’s palmitoylation-dependent localization

To identify compounds that might specifically prevent palmitoylation-dependent retrograde signaling by DLK, we expanded a high content imaging screen designed to identify small molecules that alter the palmitoylation-dependent localization of DLK-GFP in HEK cells (Fig 3A; (18)). Our expanded screen consisted of 28,400 compounds from the Maybridge and Enamine diversity sets. These libraries maximize structural diversity to increase the likelihood of identifying novel pharmacophores and are fully described under Methods. Of these 28,400 compounds, 1,723 reduced the number of cells that detectably expressed a cotransfected nuclear marker, mCherry tagged with a Nuclear Localization Sequence (mCh-NLS), by >30%. We previously used this cut-off to filter out compounds that may broadly reduce transcription and/or translation (18), so these 1723 compounds were not pursued further. For the remaining 26677 compounds, we calculated the number of DLK-GFP puncta per mCh-NLS-expressing cell (‘puncta per NLS’; P/NLS) and the average intensity of those puncta (Vesicle Average Intensity; VAI). 33 compounds increased P/NLS by a factor of >1.8 and were also not pursued further. Of the remaining 26644 compounds, 375 reduced both P/NLS and VAI by >2SD below the mean of all determinations (Fig 3B, C) and were selected for follow-up analysis.

**Figure 3:**
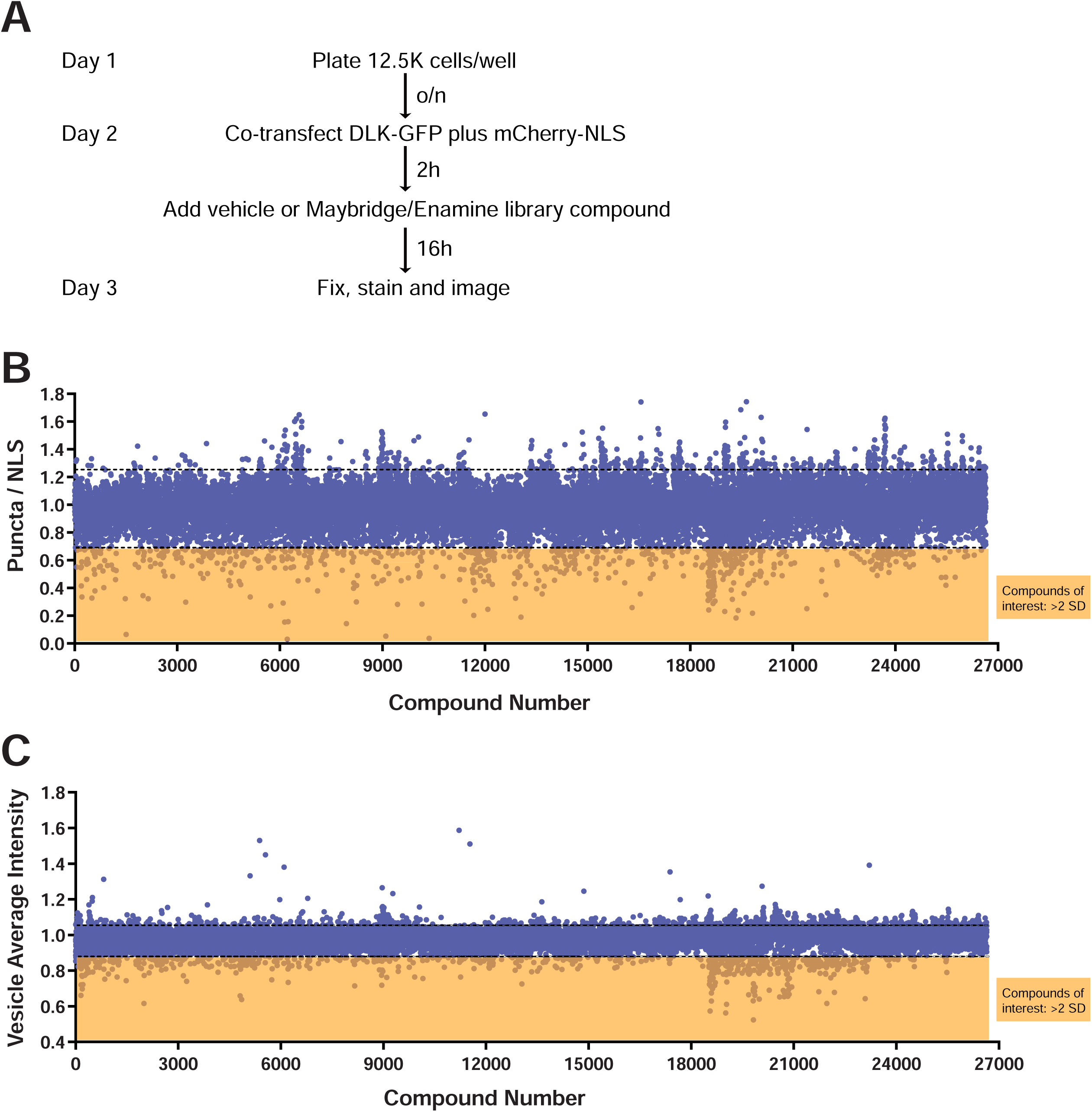
An expanded High Content Imaging screen identifies compounds that inhibit the punctate localization of DLK-GFP. ***A:*** Experimental design of the high-throughput screen to identify compounds that inhibit DLK-GFP punctate localization, adapted from (18). ***B:*** Evaluation of the effect of 26,677 compounds that passed cut-offs (of 28,400 compounds screened) from the Maybridge and Enamine Libraries™ on DLK-GFP puncta per transfected cell (puncta/NLS) and average brightness of those puncta (Vesicle Average Intensity). Black dotted lines indicate 2 standard deviations (2 SD) above and below the mean of all remaining determinations. Compounds that reduced both readouts by >2 SD were selected for further analysis.

The 375 compounds were then re-assayed in triplicate at 3 dilutions (10 μM, 3 μM, 1 μM) to confirm their effects on P/NLS and VAI and to assess dose-dependence. 33 compounds were effective in one or both readouts in this follow-up assay (Table S1). Fresh stocks of these 33 compounds (structures shown in Fig S3) were purchased for follow-up neuronal assays, in particular to determine assess their ability to inhibit palmitoyl-DLK-dependent signaling.

### Multiple hit compounds inhibit DLK-dependent c-Jun phosphorylation in trophic factor-deprived DRG neurons

We therefore asked whether compounds identified in our HEK cell screen could inhibit DLK signaling in DRG neurons subjected to TD. Following TD, DLK signals via c-Jun N-terminal kinases (JNKs), which in turn phosphorylate transcription factors including c-Jun. Phosphorylation of c-Jun (p-c-Jun) is thus a widely used readout to assess the somatic response to pro-degenerative stimuli including, but not limited to, TD (5, 8–10, 17, 26). Consistent with prior work, a DLK kinase domain inhibitor (DLKi, GNE-3511) completely blocked p-c-Jun signals induced by TD (17). Many of our hit compounds also significantly inhibited TD-induced c-Jun phosphorylation (Fig. 4A, B), suggesting that these compounds not only disrupt DLK localization in non-neuronal cells but also attenuate DLK-dependent signaling in neurons.

**Figure 4.**
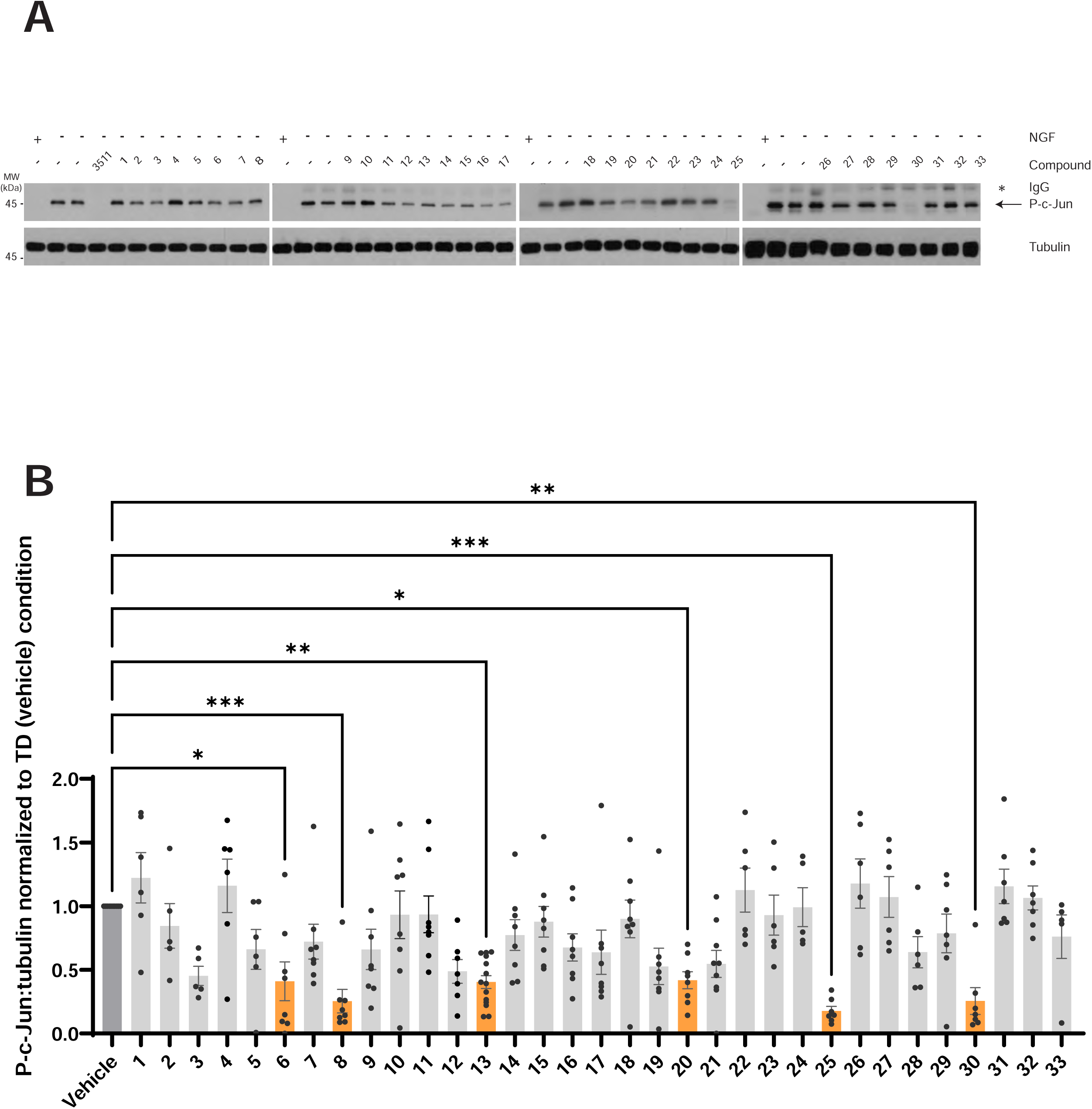
Multiple hit compounds reduce TD-induced c-Jun phosphorylation. ***A:*** Western blots of lysates from cultured DRG neurons that had been treated with vehicle (DMSO), GNE-3511 (3511) or the indicated hit compounds for 1h at 5 days in vitro (DIV 5), followed by 2.5h TD in the continued presence of the indicated compounds, or that had been left unstimulated (+ NGF). The secondary antibody used on the p-cJun blot also weakly recognizes residual anti-NGF IgG used during TD (indicated by asterisk). Subsets of compounds were assayed side-by-side, but in batches indicated by spaces between individual blots. ***B:*** Quantified data from *A*, of p-cJun:tubulin, normalized to TD (vehicle) condition. Compounds whose effect on p-c-Jun:tubulin differed significantly from TD (vehicle) condition are highlighted with orange bars. Statistical significance versus vehicle control was as follows: **6**: p = 0.021; **8**: p = 0.0007; **13**: p=0.0017; **20**: p=0.019; **25**: p=0.0002; **30**: p=0.0016, Kruskal Wallis tests with Dunn’s post hoc test for multiple comparisons.

We further investigated whether hit compounds that inhibit p-c-Jun signaling could also confer protection against degeneration in DRG neurons subjected to extended TD. Cultures subjected to TD for 45h in the presence of vehicle (DMSO) showed clear axonal blebbing/beading and cell body degeneration in bright-field images (Fig 5A, B). In contrast, both GNE-3511 and several of our hit compounds protected neuronal cell bodies and axons from TD-induced degeneration. Based on our quantitative analysis, compounds **8** and **13** were the most effective neuroprotectants and were selected for additional characterization.

**Figure 5.**
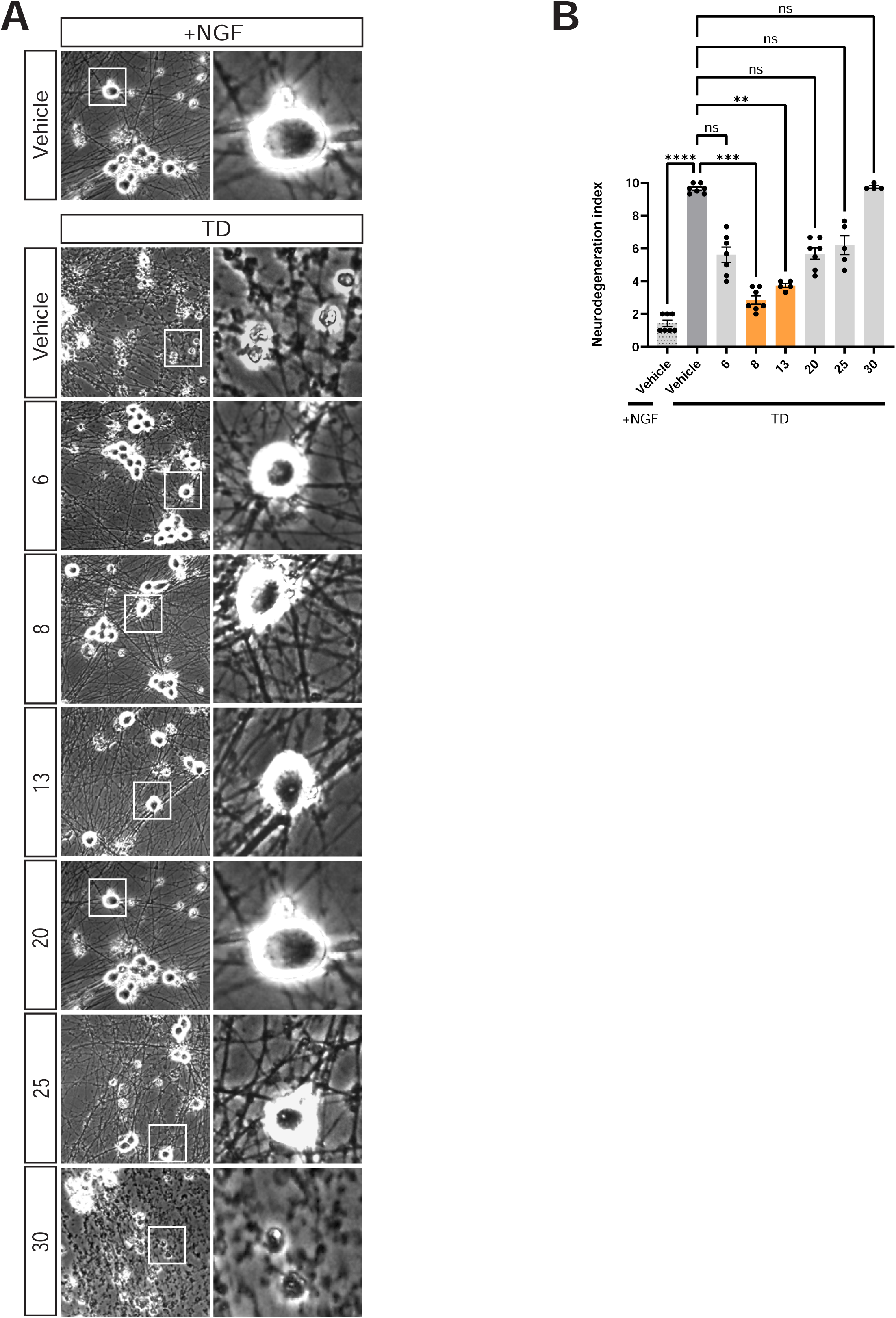
A subset of hit compounds reduce TD-induced neurodegeneration. ***A:*** Bright-field images of DRG neurons that were maintained in NGF (+NGF) or subjected to Trophic factor deprivation (TD) for 45h in the presence of vehicle (DMSO) or the indicated hit compounds that significantly reduced TD-induced c-Jun phosphorylation in Figure 4. Right-hand panels show magnified views of the boxed region in the corresponding left-hand panel. *B:* Extent of neural degeneration, quantified from images from *A*. Compounds whose effect on neurodegeneration differed significantly from vehicle control are highlighted with orange bars. Statistical significance versus vehicle control was as follows: **8**: p=0.0002; **13**: p=0.0081. TD (vehicle) condition also differed significantly from +NGF (vehicle) (p <0.0001, Kruskal Wallis tests with Dunn’s post hoc test for multiple comparisons).

### Highly neuroprotective hit compounds selectively prevent palmitoylation-dependent DLK localization and signaling after TD

Given that **8** and **13** are neuroprotective and were identified in a screen assessing DLK’s palmitoylation-dependent localization, we assessed the effect of **8** and **13** on acute, TD-induced recruitment of DLK to axonal vesicles, which is palmitoylation-dependent (Fig 2). Both **8** and **13** prevented TD-induced recruitment of DLK-myc to axonal vesicles (Fig 6A, B). Importantly, although 2BP also TD-induced recruitment of DLK to axonal vesicles, it also reduced the number of DLK-positive vesicles in healthy axons. In contrast, **8** and **13** did not affect localization of DLK to vesicles in healthy axons (Fig 6A, B). In addition, ABE assays from spot culture distal axons revealed that **8** and **13** also prevented the TD-induced increase in DLK palmitoylation (Fig 6C, D).

**Figure 6.**
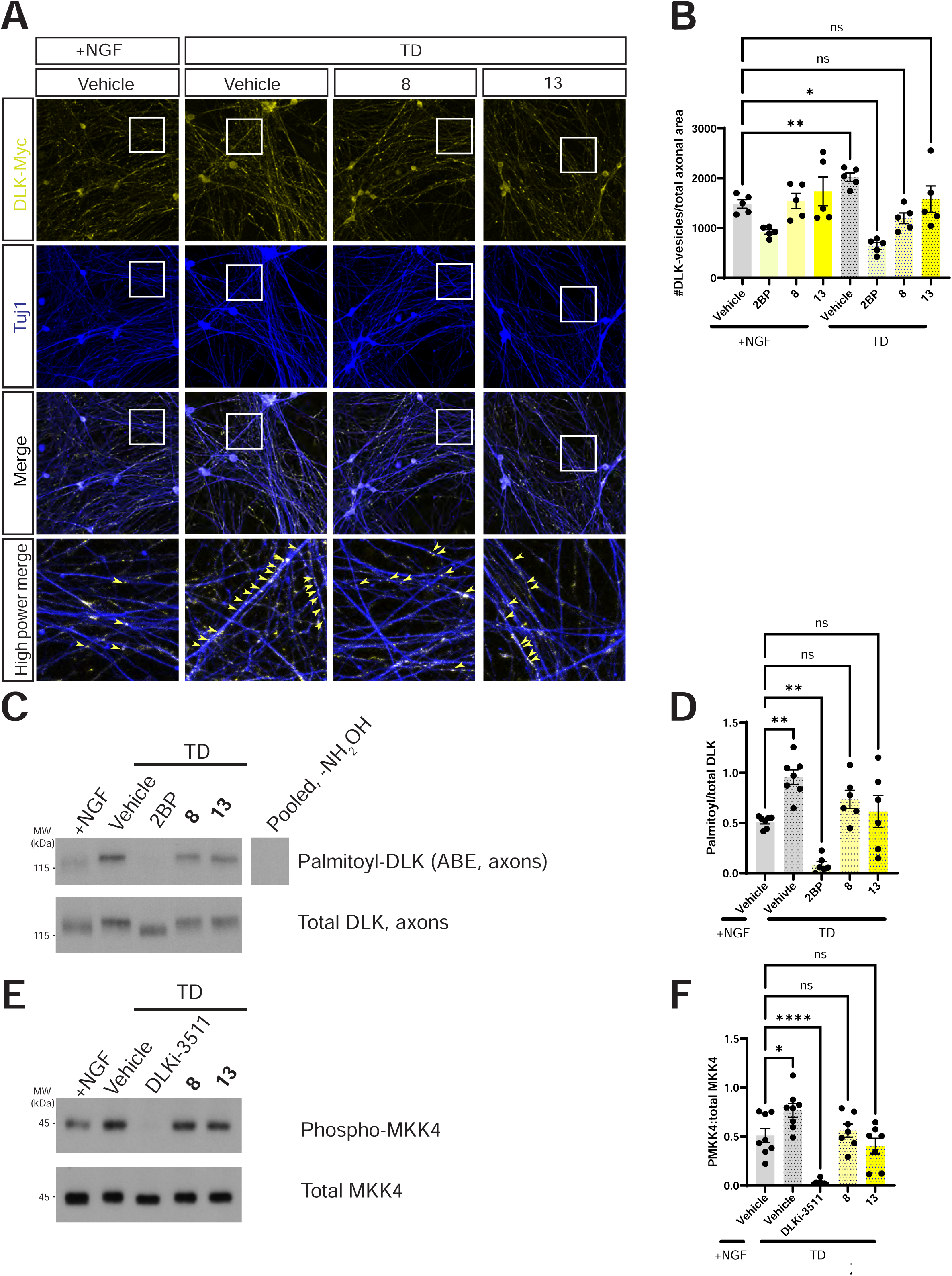
Neuroprotective hit compounds prevent TD-induced DLK recruitment to axonal vesicles and palmitoylation, and activation of DLK’s downstream target MKK4. *A:* Cultured DRG neurons were infected to express myc-tagged wild type DLK (wtDLK-myc) and left untreated or were subjected to TD in the presence of the indicated compounds or DMSO vehicle, prior to fixation and immunostaining with the indicated antibodies. The bottom row of images shows a magnified view of the boxed region in upper panels. *B:* Quantified data from *A* confirm that TD-induced increases in axonal DLK puncta (presumptive vesicles (24)) are prevented by **8** and **13**. Data for +NGF (vehicle), TD (vehicle) and TD (2BP) conditions are replotted from Fig 2B. *; p=0.0483; **; p=0.0016, ANOVA, Dunnett’s post hoc test. *C:* Western blots of ABE (palmitoyl-) fractions of DRG axonal lysates that had been treated as indicated prior to lysis. *D:* Quantified data from *C* confirm that **8** and **13** prevent the TD-induced increase in axonal DLK palmitoylation. Data for +NGF (vehicle), TD (vehicle) and TD (2BP) conditions are replotted from Fig 2D. +NGF (vehicle) versus TD (vehicle): **; p=0.0026; +NGF (vehicle) versus TD (2BP): **; p=0.0054, ANOVA, Dunnett’s post hoc test. The righthand lane is a side-by-side exposure from a parallel control sample omitting the key ABE reagent NH_2_OH, run on the same gel, with intervening spacer lanes cropped. *E:* Western blots, detected with the indicated antibodies, of DRG lysates that had been treated as indicated prior to lysis. *F:* Quantified data from *E* reveal that TD significantly increases pMKK4:total MKK4 (+NGF (Vehicle) vs. TD (Vehicle); p =0.0219) while DLK inhibitor GNE-3511 (DLKi-3511) significantly reduces it (+NGF (Vehicle) vs. TD (DLKi-3511): p<0.0001) but TD has no effect on pMKK4:total MKK4 in the presence of **8** or **13** (ns; non-significant). ANOVA, Dunnett’s post hoc test.

The effects of **8** and **13** on DLK’s axonal palmitoylation and localization suggested that they might also prevent TD-induced signaling by DLK. Indeed, when we assessed DLK’s direct substrate, MKK4, we found that **8** and **13** both prevented TD-induced increases MKK4 phosphorylation (pMKK4) (Fig 6E, F). In contrast, neither **8** nor **13** phenocopied the mislocalization of axonal NF-200, Tuj1, DLK and VAMP2 seen with GNE-3511 (Fig7A-F). Together, these findings suggest that **8** and **13** selectively inhibit pro-degenerative retrograde axonal signaling by DLK, but minimally affect roles of DLK in healthy axons.

### Protective compounds identified in primary neurons blunt prodegenerative retrograde signaling in vivo

Finally, we assessed the translational potential of **8** and **13**. RGC degeneration after optic nerve crush (ONC) is highly dependent on (palmitoyl)-DLK (6, 7, 9, 27) and local, intravitreal delivery of compounds to the eye could circumvent potential issues with systemic instability. Because intravitreally injected compounds can rapidly drain from the eye (28, 29), we isolated retinas 15h post-ONC, after first confirming that ONC-induced c-Jun phosphorylation is robustly observed at this time point (Fig S4). Importantly, intravitreal injection of either **8** or **13** reduced DLK-dependent responses of RGCs (ONC-induced c-Jun phosphorylation) to a similar extent as GNE-3511 (Fig 8A, B; RGCs identified with the marker Brn3A). These findings suggest that **8** and **13** inhibit DLK-dependent axon-to-soma signaling in vivo. These results reveal a novel therapeutic approach that may more selectively block pro-degenerative retrograde signaling by DLK without the side effects of globally inhibiting all pools of this kinase.

## Discussion

Given the numerous preclinical studies that identified DLK as a promising therapeutic target (6, 7, 11, 12, 30), the number of patients who experienced sensory neuropathy and other adverse events in a recent clinical trial of a DLK kinase domain inhibitor was disappointing (16). The unexpected finding of elevated neurofilament levels in patient plasma during this trial indicate that DLK plays a previously unappreciated role in maintaining the integrity of the neuronal cytoskeleton (16). The association between DLK inhibitor treatment and sensory neuropathy symptoms further suggested that DLK might directly regulate the cytoskeleton of DRG sensory neurons (16). One major finding from our study is the direct experimental support for this latter hypothesis, as we reveal that acute DLK inhibition disrupts axonal neurofilament and tubulin distribution and causes an accumulation of axonal vesicles in DRG neurons (Fig 1). These striking findings are being further investigated and will be reported in more detail in future studies, but one initial conclusion, consistent with (16), is that global inhibition of DLK kinase activity induces unintended side effects in neuronal axons.

Nonetheless, targeting other features of DLK, including specific interactions and/or post-translational modifications might still be of therapeutic benefit. In this study, we therefore focused on the regulation of DLK by palmitoylation, a modification that is critical for both axonal localization and signaling by DLK (8, 9, 17, 31). These conclusions were in part based on experiments in which loss of the protein acyltransferase (PAT) ZDHHC17, which plays an evolutionarily conserved role in controlling DLK palmitoylation and subsequent targeting to neuronal axons, was also found to block DLK-dependent retrograde signaling (9). It was therefore intriguing that a subsequent study additionally suggested that a distal axonal pool of DLK is acutely palmitoylated after TD (24). Given that ZDHHC17 is a Golgi-localized PAT (9, 32, 33), this finding might seem at odds with our prior results. Importantly, though, palmitoylation is a reversible modification. A plausible model to explain these findings is thus that ZDHHC17-dependent palmitoylation on the Golgi facilitates vesicle-based transport of DLK to distal axons, where a subset of DLK molecules is depalmitoylated and is then acutely re-palmitoylated after TD, likely by a PAT distinct from ZDHHC17. Control of distinct subcellular pools of a given palmitoyl-protein by different PATs is not without precedent (34) and raised the possibility of specifically targeting this stimulus-dependent post-translational modification event for therapeutic benefit.

While striking on a cell biological level, the functional importance of acute axonal palmitoylation of DLK (24) for retrograde pro-degenerative signaling had not been previously investigated. Moreover, although our own studies had revealed a critical functional role for DLK palmitoylation (8, 9), these studies had employed a DLK mutant that is never palmitoylated, and thus could not distinguish the importance of basal, Golgi-localized palmitoylation from that of any stimulus-dependent axonal palmitoylation of DLK. In this study we confirm that acute, TD-induced palmitoylation of DLK indeed occurs and is concomitant with DLK recruitment to axonal vesicles (Fig 2A-D). Importantly, we further reveal that axonal palmitoylation is critical for retrograde DLK-dependent signaling (Fig 2E, F), supporting the notion that this event might be pharmacologically targeted to prevent DLK-dependent retrograde signaling, potentially without affecting other roles of DLK.

Although inhibiting stimulus-dependent palmitoylation of DLK would appear to be the most selective approach, our high content screen could potentially identify hits that predominantly affect basal DLK palmitoylation, stimulus-dependent palmitoylation, or both. Indeed, given that the majority of DLK is Golgi-localized in HEK293T cells (9, 18), the screen likely favors identification of the first of these classes of compounds. It is thus intriguing that **8** and **13** were far from the most potent disruptors of DLK punctate localization in our primary screen (Table S1) yet were the most effective inhibitors of palmitoylation-dependent DLK-dependent signaling and degeneration in neuronal assays. These findings are consistent with a model in which the primary mechanism of action of **8** and **13** is to selectively disrupt stimulus-dependent palmitoylation of DLK. This conclusion is further supported by our finding that neither of these compounds disrupts DLK’s axonal vesicular targeting in healthy axons (Fig 6B).

Importantly, while this study identifies neuroprotective molecules with a distinct mechanism of action from DLK kinase domain inhibitors, it is likely premature to suggest that **8** or **13** themselves should be pursued as therapeutic agents. In particular, we recognize that the block of ONC-induced p-c-Jun by **8** and **13**, while promising for potential in vivo applications, is only partial (Fig 8A, B). There may, however, be multiple reasons for this incomplete inhibition. For example, the extent to which these compounds can diffuse from the RGC soma into the proximal axon, where stimulus-dependent palmitoylation is most likely to occur in vivo, is unclear and may limit their therapeutic efficacy. A second issue is that small molecules are rapidly cleared from the mouse eye after intravitreal injection (28, 29) which likely limits the effective concentration of **8** and **13** in the hours after ONC. However, controlled release strategies using intravitreal implants e.g. (35, 36) could be an attractive approach for diseases of the visual system in which DLK is implicated (7). Importantly, though, our findings suggest that similar strategies, using more expansive libraries and/or involving combinatorial chemistry to identify more potent analogs of **8** and/or **13**, could identify compounds that could indeed act as potent in vivo neuroprotectants. Our results thus reveal a highly promising alternative approach to target specific pools of palmitoyl-DLK, which could serve as a novel therapeutic strategy that circumvents issues associated with global DLK inhibition.

## Experimental Procedures

### Molecular biology, cDNA constructs

DLK-GFP and pmCherry-NLS (the latter kindly provided by Martin Offterdinger (Addgene Plasmid #39319)) were previously described (18). The lentiviral vector FEW-DLK-myc was previously described (8). For this study the EF1 alpha promoter in this vector was replaced with a human synapsin promoter by standard subcloning, generating FSW-DLK-myc, which ensures neuron-specific DLK expression.

### Screening libraries and high content imaging

This study employed a small molecule diversity-based screening library, selected from the Maybridge and Enamine Screening Collections. This combined library maximizes structural diversity in order to increase the chances of identifying novel pharmacophores. The library obeys Lipinski rules, with all logP values < 5 (average logP value = 3.2), < 5 H-bond donors, < 10 H-bond acceptors, number of rotatable bonds < 8 (average # of rotatable bonds < 5) and molecular weights < 500 (average MW = 325). All potentially reactive molecules have been removed. Structural integrity of the library members was originally confirmed by the vendors using ^1^H-NMR and LC/MS and reconfirmed by Temple’s Moulder Center using LC/MS. Samples were originally purchased as powders and formulated into 10 mM DMSO stock solutions at the Moulder Center. 20,000 Maybridge compounds and 8,400 Enamine compounds were screened for this study. For follow-up assays in neurons, individual compounds were repurchased from Maybridge and Enamine as solids and were reconstituted as 10mM (for cultured neuron experiments) or 100mM (for in vivo experiments) 1000x stocks in DMSO.

### Chemicals

2-Bromopalmitate (2BP) and S-Methyl methanethiosulfonate (MMTS) were from MilliporeSigma. HPDP Biotin was from Soltec Ventures. Microcystin-LR and DLK inhibitor GNE-3511 were from Cayman Chemicals. All other chemicals were from Thermofisher Scientific and were of the highest reagent grade.

### Antibodies

The following primary antibodies, raised in the indicated species, were used: anti-NGF (sheep, CedarLane, #CLMCNET-031); phospho–c-Jun (Ser^63^) (rabbit, Cell Signaling Technology, #91952); DLK/MAP3K12 (rabbit, Genetex, #GTX124127); GFP (rabbit; Life Technologies, #A11122); β3 tubulin (mouse, BioLegend, TUJ1, #MMS-435P), myc (rabbit, Cell Signaling Technology #2278), alpha-tubulin (mouse, Cell Signaling Technology, #3873), phospho-MKK4 (rabbit, Cell Signaling Technology, #4514), pan-MKK4 (rabbit, Cell Signaling Technology, #9152), NF-200 (mouse, Sigma #NO-142), VAMP2 (mouse, Synaptic Systems, #104211SY), Brn3a (mouse, Millipore Sigma, # MAB1585): .

### Cell transfection

HEK293T cells were transfected using a calcium phosphate-based method as described previously (37).

### High Content Screening (HCS) assay

HCS assay was performed essentially as previously described (18). Briefly, HEK293T cells were seeded in poly-lysine coated 96 well plates (Greiner Bio-One, black walled chimney-wells), transfected as above and treated with 2BP (10 μM final concentration), library compounds or DMSO vehicle control at 2 h post-transfection. Maybridge or Enamine library compounds were spotted onto 96 well plates at 10 mM in DMSO and resuspended in 200 µL pre-warmed DMEM. 40 µL of diluted compound was then added to cells in 160 µL of DMEM (containing glutamax, 10% FBS and antibiotics). Cells were returned to a tissue culture incubator for a further 14 h at 37 °C. Subsequently, medium was removed and cells were fixed in 4% PFA (1x PBS) for 20 mins at RT, washed once with PBS and stained with 300 nM DAPI for 5 mins at RT, followed by 2 washes of PBS.

### High Content Imaging

High Content imaging was performed as in (18) using an ImageXpress micro high content imaging system (Molecular Devices, Downingtown, PA) driven by MetaXpress software. Six images per well were acquired in each of three channels (DAPI, FITC, TRITC) at 10X magnification in an unbiased fashion.

### Analysis of high content imaging data

Images were analyzed as in (18) using the MetaXpress ‘Multiwavelength Scoring’ (for mCherry-NLS signals) and ‘Transfluor’ modules (for DLK-GFP signals). Data were exported to a spreadsheet using the AcuityXpress software package (Molecular Devices). Three metrics were used: DLK puncta (“Total Puncta Count” option, from DLK-GFP signal), DLK vesicle average intensity (VAI; the intensity of the punctate DLK-GFP signal) and total number of transfected cells (from mCherry-NLS signal). The first and last of these metrics were combined to calculate DLK-GFP Puncta per NLS (P/NLS). Compounds that reduced P/NLS by greater than 30% of the average of vehicle-treated controls for each day were excluded from analysis due to likely cytotoxicity and/or broad effects on transcription, translation or protein stability. Compounds that increased P/NLS by >1.8 fold were also excluded from further analysis. Compounds that reduced DLK-GFP P/NLS and VAI by 2 times the standard deviation of the mean of all remaining determinations were considered “Hits”. This calculation was performed on a running basis in order to follow up initial hits while the primary screen was ongoing. The 2SD cut-offs plotted in Figure 3 represent the final calculated SD values after all compounds had been screened. Follow-up assays of hit compounds were performed as above except that 10mM stocks of compound were manually diluted to 3mM and 1mM in DMSO and were added to triplicate wells of a 96-well plate containing transfected cells (10μM, 3μM, 1μM final concentration).

### DRG conventional, ‘spot’ and microfluidic cultures

Primary dorsal root ganglion (DRG) neurons were isolated from embryonic day 16 (E16) rat embryos. All procedures followed the National Institutes of Health guidelines and were approved by the Institutional Animal Care and Use Committee (IACUC) at Temple University. Conventional ‘mass’ cultures were plated on either tissue culture plastic or glass coverslips pre-coated with poly-lysine and laminin, as previously described (8). ‘Spot’ cultures were plated on tissue culture plastic pre-coated with poly-lysine and laminin similar to (38). Microfluidic cultures were prepared as in (8, 39), based on a previously described design (40).

### Trophic deprivation assay in conventional cultures

Conventional DRG cultures, prepared as above, were treated at 5 days in vitro (DIV 5) with either DMSO or 10 µM of hit compounds for 1 hour prior to trophic factor deprivation (TD). For all TD experiments, NGF-containing medium was replaced with fresh Neurobasal medium lacking NGF but containing B27 supplement plus sheep anti-NGF antibody (25 μg/ml), in the continued presence of drug or vehicle. For biochemical experiments, cells were lysed 2.5h post-TD in SDS-PAGE loading buffer and then processed for subsequent SDS-PAGE and immunoblotting analysis. For immunocytochemistry, cultures were fixed 3h post-TD. For assays of neurodegeneration, cultures were imaged live 45h post-TD using an Olympus CKX41 inverted microscope with 20X, 0.4 NA objective. The extent of degeneration was quantified manually on a 10-point scale, similar to prior studies (9, 41), by an experimenter blinded to the treatment condition. Importantly, TD-induced responses in conventional cultures require the retrograde motor protein dynein (9) so this assay likely directly assesses retrograde axonal signaling, a process known to require palmitoyl-DLK (8, 9).

### Retrograde signaling assay in microfluidic cultures

E16 DRG neurons were plated in microfluidic chambers as in (8, 39), based on a previously described design (40). Selective TD of distal axons was performed on DIV8-10 and cultures were processed for immunocytochemistry as in (17).

### Acyl biotin exchange (ABE) palmitoylation assay from axonal fractions of DRG neurons

E16 DRG neurons were plated as spot cultures and at DIV9 were pre-treated for 1h with either vehicle, 2BP, or hit compounds. TD was then performed in the continued presence of hit compounds or vehicle. 2h later, a biopsy punch was used to separate the axon and soma fractions, both of which were immediately denatured in ABE lysis buffer (37). ABE was performed essentially as in (37), except that after blocking with MMTS, cultures were mixed with 1 ml of 0.08% (v/v) carrier protein (heat inactivated FBS that had itself been blocked with MMTS). Subsequent acetone precipitation, Biotin-HPDP treatment and capture on neutravidin beads and elution were performed as previously described (37).

### Lentiviral preparation and infection of DRG cultures

VSV-G pseudotyped lentivirus was prepared as previously described (37). The minimum amount of virus needed for subsequent immunocytochemical detection of virally expressed wtDLK-myc was determined in pilot studies and that amount was then infected on the second day in vitro (DIV2). On DIV5, infected cultures were pre-treated with DMSO vehicle or inhibitors for 1h, subjected to TD for 3h in the continued presence of vehicle or inhibitors, and subsequently fixed and immunostained.

### Immunocytochemistry of cultured DRG neurons

Immunostaining of dissociated DRG neurons cultured on coverslips was performed essentially as described (17). Briefly, coverslips were rinsed once with 1× recording buffer [25 mM Hepes (pH 7.4), 120 mM NaCl, 5 mM KCl, 2 mM CaCl2, 1 mM MgCl2, and 30 mM glucose] and fixed for 10 min in 4% PFA/sucrose diluted in PBS at room temperature. Samples were permeabilized in PBS containing 0.25% (w/v) Triton X-100 for 10 min at 4°C, blocked with PBS containing 10% (v/v) normal goat serum (SouthernBiotech, 0060-01) for 1 hour, and incubated in primary antibodies overnight at 4°C in blocking solution. After three PBS washes, coverslips were incubated for 1 hour at room temperature with Alexa Dye–conjugated fluorescent secondary antibodies diluted in blocking solution, prior to four final PBS washes and mounting using FluorSave reagent (MilliporeSigma). Images shown in Figure 1 and Figure 7 were acquired using a Nikon C2 inverted confocal microscope with an oil immersion objective (60×, 1.4 NA). Acquisition parameters (laser power, gain and offset) were kept constant between all conditions. Maximum intensity projections were generated using NIS Elements software and exported to ImageJ/Fiji for analysis. Images in Figure2A, Figure 6 and Figure 8 were acquired using a Leica SP8 confocal microscope with oil immersion objective. Images in Figure 2E were acquired using a Nikon 80i fluorescence microscope with a 10×, 0.3 NA objective.

**Figure 7:**
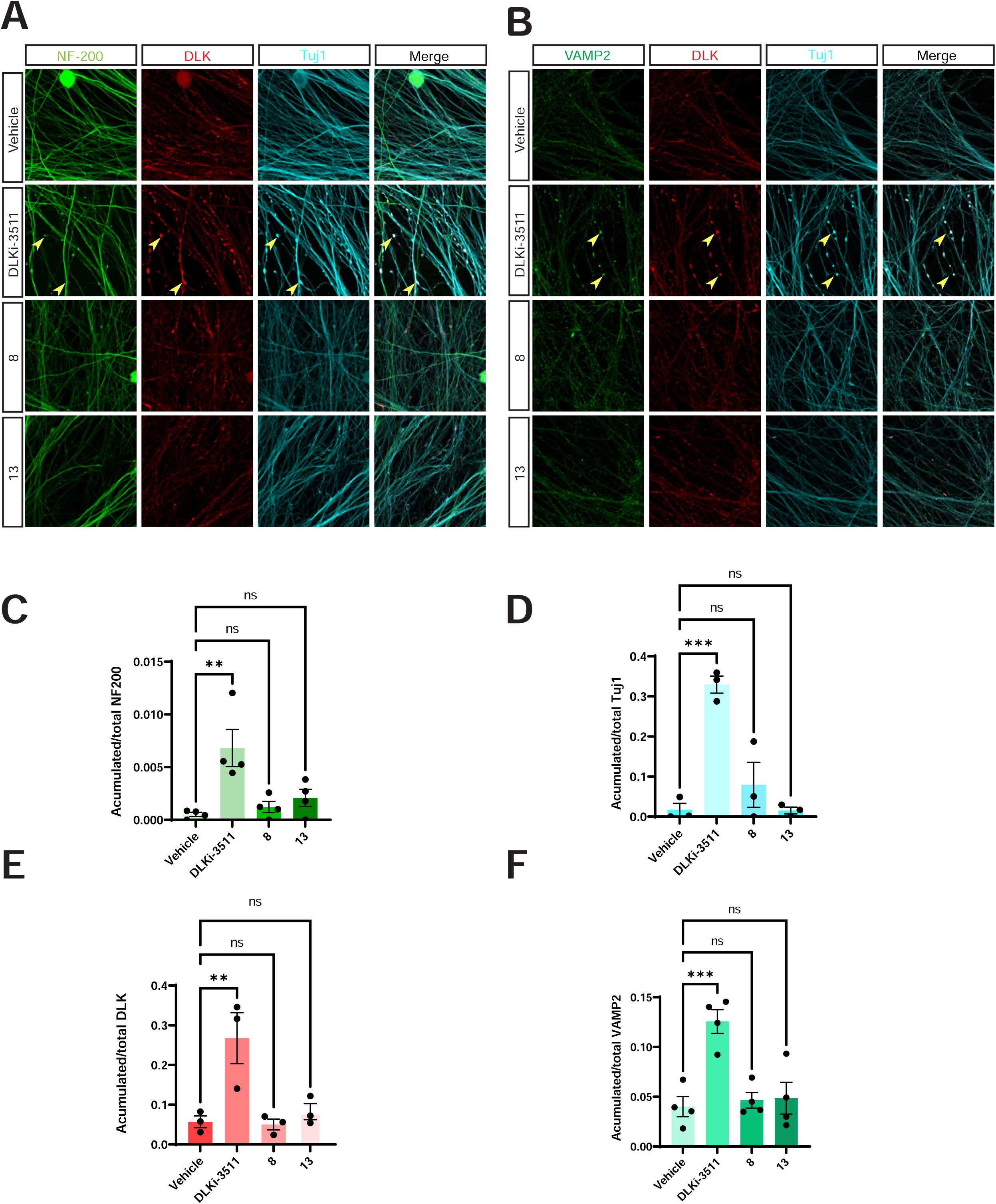
Neuroprotective hit compounds do not phenocopy the effect of a DLK kinase domain inhibitor on axonal integrity. ***A:*** Images of cultured DRG neurons treated with DMSO vehicle, 500 nM DLKi-3511 or 10 μM of the indicated compounds for 4 hours and fixed and stained with the indicated antibodies. ***B:*** As *A*, except that neurons were fixed to detect VAMP2 rather than NF-200. Quantified data from *A* and *B* confirm that **8** and **13** do not cause disruptions of NF-200 (***7C***: **; p=0.0022) or Tuj1 (***7D***: ***; p=0.0003), nor do they induce accumulation of DLK (***7E***: **; p=0.0072) or VAMP2 (***7F***: ***; p=0.0007) in axons. ANOVA with Dunnett’s post hoc test for all panels *C-F*.

**Figure 8:**
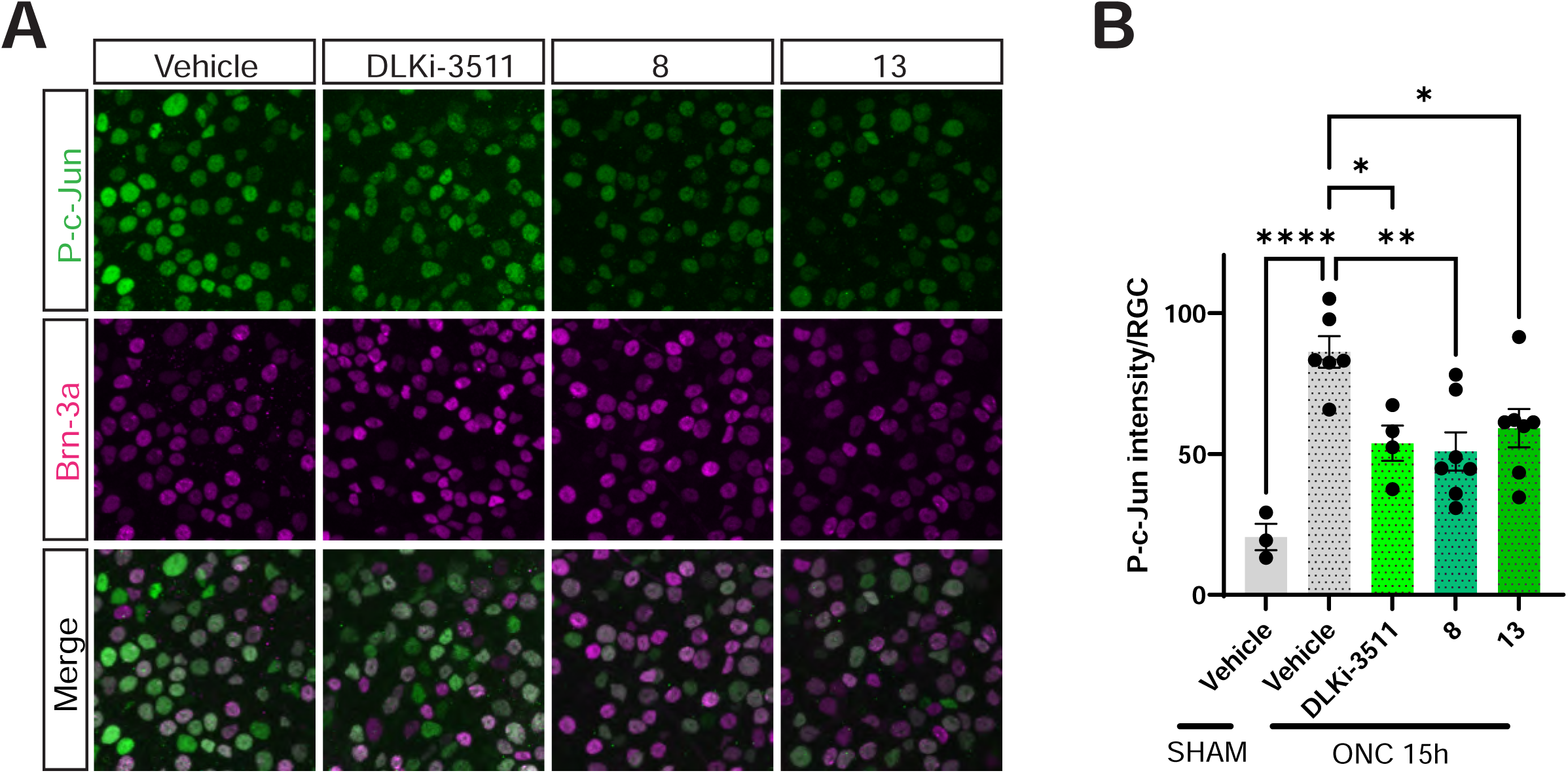
Hit compounds reduce palmitoyl-DLK-dependent pro-degenerative retrograde signaling to a similar extent as a DLK kinase domain inhibitor in vivo. *A:* Images of retinas of mice that had been subjected to optic nerve crush (ONC), intravitreally injected with the indicated compounds or DMSO vehicle and then perfused, fixed and immunostained with the indicated antibodies 15h post-ONC. *B:* Quantified data from *A* reveal that DLKi-3511, **8** and **13** inhibit ONC-induced c-Jun phosphorylation. ONC (vehicle) versus ONC (DLKi-3511); *; p =0.0144;), ONC (vehicle) versus ONC (**8**): **; p =0.0019; ONC (vehicle) versus ONC (**13**): *; p =0.0181. Sham (vehicle) and ONC (vehicle) conditions also differ significantly (****; p<0.0001), ANOVA, Dunnett’s post hoc test.

### Analysis of images from cultured DRG neurons

Images of endogenous DLK, VAMP2, NF200 and Tuj1 were auto-thresholded in NIH ImageJ/Fiji. The ‘analyze particles’ algorithm was then used to detect axonal accumulations of size 200-1500 pixels and with circularity 0.2-1.0 (DLK, VAMP2) or 0.4-1.0 (NF200, Tuj1) The percentage of axonal area occupied by these accumulations (total particle area) was then expressed as a fraction of the total axonal area as defined by each channel.

For analysis of DLK-myc images, DRG cell bodies were manually identified and cleared from each image in ImageJ/Fiji, to leave only axonal signals. Images were auto-thresholded and puncta of size 4.0-10.0 pixels and circularity 0.70-1.00 were counted. The number of puncta was normalized to the fraction of the total area of the field occupied by axons, as determined by the Tuj1 channel.

### Intravitreal injection and Optic nerve crush

Six-week-old mice were anesthetized with 0.01 mg of xylazine and 0.08 mg of ketamine per gram of body weight prior to intravitreal injection of the right eye with 2 μl of either DLKi-3511 (20 μM), **8** or **13** (100 μM),) or an equivalent volume of DMSO vehicle, all diluted in 0.5× sterile PBS containing 1% (w/v) sorbitol and 5% (v/v) glycerol. The injection needle was carefully inserted behind the orthoptic lens and into the vitreous chamber to avoid damaging the lens. The left eye was left untreated. Immediately after intravitreal injection, while mice were still anesthetized, optic nerve crush (ONC) was performed as previously described (9). In the sham group, the right eye was intravitreally injected with 2 μl of DMSO/buffer as above but was not subjected to ONC. Fifteen hours after ONC or sham injury, mice were transcardially perfused with 4% paraformaldehyde (PFA) in 1× phosphate-buffered saline (PBS). Eyeballs were postfixed for an additional two hours before dissection of the retinas. Whole-mount retinal staining was performed as in (9) using DAPI and antibodies raised against p-c-Jun, Tuj1 and the RGC marker Brn3A.

### Image acquisition and analysis of ONC samples

DAPI, p-c-Jun and Brn3A signals from flat-mounted retinas were acquired using a Leica SP8 confocal microscope with a 40x oil immersion objective. Maximum intensity projections were generated from confocal z-stacks and individual RGC nuclei were then identified by creating a mask of the overlapping DAPI and Brn3a signals in Fiji/ImageJ, using the Image Calculator ‘AND’ function. This nuclear mask was then used to calculate the average intensity of the phospho–c-Jun signal in each individual RGC nucleus. For each experimental eye, phospho–c-Jun intensity within each RGC was calculated for three separate fields of view, which were then averaged to produce a single ‘n’ per eye.

### Experimental replicates and statistical analysis

For all experiments using cultured neurons, ‘n’ reflects the indicated number of cultures, each from different dissections, indicated as individual data points in each Figure. In some cases, replicate determinations from a single dissection were performed side-by-side and averaged to give a single biological ‘n’. For immunocytochemical studies in neurons, an average of three to four images per coverslip were acquired and analyzed to produce one biological replicate for plotting.

Statistical analysis was performed using GraphPad Prism. Two-group comparisons of normally distributed variables were conducted using a t-test, applying Welch’s correction for unequal variances or sample sizes. ANOVA was used for multi-condition experiments, with Dunnett’s post hoc test for comparisons against an indicated control condition. For non-normally distributed continuous variables, the Mann-Whitney U test was used for two-group comparisons, and the Kruskal-Wallis test with Dunn’s test for multiple comparisons for more than two groups.

## Supporting information

Supplementary Material Zhang

## Acknowledgements

We thank Natasha Hesketh for assistance with DRG cultures and invaluable suggestions, Dr. Azita Minaei (Qazvin University of Medical Sciences, Iran) for insightful comments on the manuscript, Luiselys Hernandez for help with characterization of hit compounds in neurons and Drs Marlene Jacobson and Dale Martin for initial optimization of DLK screening. Supported by grants from NIH (R01 NS094402 and R21 EY029386, both to G.M.T.; NCI R50 CA211479 to M.B.E. and NCI Core Grant P30 CA006927 to Fox Chase Cancer Center) and by Shriners’ Childrens (#85190 PHI and #87400 PHI, both to G.M.T.).

