## Supplementary Material Zhang for "Novel inhibitors of acute, axonal DLK palmitoylation are neuroprotective and avoid the deleterious side effects of cell-wide DLK inhibition"

**Supplementary Table:**

**Table S1: Data for 33 compounds from primary screen that reduced DLK-GFP puncta number (puncta per NLS; P/NLS) or intensity (vesicle average intensity; VAI) in a follow-up triplicate assay.** All compounds from primary screen that reduced both P/NLS and VAI below the 2SD cut-off were re-assayed in triplicate at the indicated concentrations using the HEK 293T cell assay shown in Figure 2. Data for 33 compounds that reduced (i) P/NLS by >30% and/or (ii) VAI by >10% in this follow-up assay are plotted here. None of these compounds reduced mCherry NLS count by >30% in the primary screen, although a subset of compounds did so in this follow-up screen. However, all compounds that met criteria (i) and/or (ii) were assayed in neurons (Fig 4).

| | | Puncta/NLS (P/NLS) | | | | | | Vesicle Average Intensity (VAI) | | | | | | mCherry NLS | | | | | |
|---|---|---|---|---|---|---|---|---|---|---|---|---|---|---|---|---|---|---|---|
| | | 10uM | | 3uM | | 1uM | | 10uM | | 3uM | | 1uM | | 10uM | | 3uM | | 1uM | |
| Hit No | **Maybridge CAT_No** | AVE | % | AVE | % | AVE | % | AVE | % | AVE | % | AVE | % | AVE | % | AVE | % | AVE | % |
| **1** | BTB00713SC | 1.06915 | **30.9** | 1.51491 | 2.1 | 1.84922 | -19.5 | 412.482 | **11.8** | 447.155 | 4.4 | 463.591 | 0.9 | 327.556 | 1.8 | 343.778 | -3.0 | 317.611 | 4.8 |
| **2** | BTB07602SC | 0.49728 | **43.9** | 0.82423 | 6.9 | 1.33725 | -51.0 | 379.586 | **17.3** | 423.236 | 7.8 | 461.226 | -0.4 | 569.833 | -18.0 | 592.333 | -22.6 | 457.5 | 5.3 |
| **3** | CD01955SC | 0.88549 | **42.8** | 1.45184 | 6.2 | 1.62339 | -4.9 | 414.405 | **11.4** | 452.981 | 3.2 | 461.601 | 1.3 | 349 | -4.6 | 347 | -4.0 | 346.722 | -3.9 |
| **4** | CD02036SC | 0.74653 | **47.5** | 1.34663 | 5.4 | 1.62787 | -14.4 | 392.793 | **13.0** | 441.055 | 2.3 | 447.077 | 0.9 | 397.278 | -30.5 | 358.333 | -17.7 | 305.556 | -0.3 |
| **5** | HTS01118SC | 0.3442 | **52.8** | 0.6201 | 14.9 | 0.52028 | 28.6 | 416.815 | **10.7** | 448.076 | 4.0 | 456.795 | 2.2 | 532.778 | 2.5 | 496.833 | 9.1 | 544 | 0.5 |
| **6** | HTS04553SC | 0.56684 | 22.3 | 0.8891 | -21.9 | 0.82925 | -13.7 | 416.925 | **10.7** | 459.938 | 1.5 | 472.364 | -1.2 | 586.444 | -7.3 | 500.111 | 8.5 | 547.778 | -0.2 |
| **7** | HTS05834SC | 0.57781 | **33.6** | 1.04635 | -20.2 | 1.17442 | -34.9 | 384.126 | **16.4** | 442.331 | 3.7 | 464.852 | -1.2 | 386.111 | 7.5 | 394.722 | 5.4 | 391.722 | 6.2 |
| **8** | HTS06895SC | 0.33305 | **36.6** | 0.42242 | 19.6 | 0.43848 | 16.6 | 338.79 | **13.7** | 375.844 | 4.3 | 379.905 | 3.3 | 141.222 | **32.2** | 174.445 | 16.3 | 167.611 | 19.6 |
| **9** | HTS09450SC | 0.30437 | **47.9** | 0.42665 | **27.0** | 0.43362 | **25.8** | 357.98 | **13.8** | 379.69 | 8.6 | 418.847 | -0.9 | 173.778 | 28.0 | 161.722 | 33.0 | 136 | 43.7 |
| **10** | RH01693SC | 0.13843 | **75.5** | 0.47056 | 16.6 | 0.58938 | -4.4 | 347.969 | **14.0** | 386.824 | 4.4 | 391.114 | 3.4 | 195.444 | **31.1** | 288.611 | -1.7 | 363.722 | -28.2 |
| **11** | RJC01887SC | 0.31679 | **39.7** | 0.47291 | 10.0 | 0.47338 | 9.9 | 343.22 | **12.6** | 366.428 | 6.7 | 378.858 | 3.5 | 238.834 | -14.6 | 277.556 | -33.2 | 300.055 | -44.0 |
| **12** | RJC03231SC | 0.27939 | **51.4** | 0.51674 | 10.2 | 0.56742 | 1.4 | 342.367 | **15.0** | 379.058 | 5.9 | 396.732 | 1.5 | 260.778 | -19.8 | 278.111 | -27.8 | 303.889 | -39.6 |
| **13** | RJF01488SC | 0.25292 | **57.4** | 0.42584 | **28.2** | 0.54882 | 7.5 | 314.532 | **24.3** | 367.359 | **11.6** | 385.751 | 7.2 | 183.667 | 16.5 | 247.889 | -12.8 | 298.611 | -35.8 |
| **14** | S09668SC | 0.22553 | **53.1** | 0.30046 | **37.6** | 0.42357 | 12.0 | 363.298 | **12.3** | 382.363 | 7.7 | 413.192 | 0.2 | 108.666 | **32.8** | 151.278 | 6.5 | 226.556 | -40.0 |
| **15** | SEW02506SC | 0.15489 | **70.5** | 0.48436 | 7.9 | 0.46333 | 11.9 | 295.78 | **24.7** | 380.431 | 3.1 | 380.918 | 3.0 | 140.167 | **32.7** | 290.667 | -39.5 | 254.778 | -22.3 |
| **16** | SEW04994SC | 0.31607 | **45.1** | 0.50192 | 12.8 | 0.58684 | -2.0 | 339.02 | **15.8** | 379.611 | 5.7 | 381.644 | 5.2 | 143.167 | **34.2** | 237.056 | -8.9 | 266.167 | -22.3 |
| **17** | TL00098SC | 0.24659 | **56.3** | 0.32358 | **42.7** | 0.39694 | **29.7** | 345.797 | **14.5** | 359.271 | **11.2** | 363.715 | **10.1** | 239.889 | 15.5 | 241.833 | 14.8 | 240.666 | 15.2 |
| **18** | BTB10119SC | 0.13547 | **68.9** | 0.20537 | **52.9** | 0.29332 | **32.7** | 331.408 | **12.9** | 353.035 | 7.2 | 358.024 | 5.9 | 175.667 | 18.5 | 216.167 | -0.3 | 274.778 | -27.5 |
| **19** | DP01302SC | 0.32674 | **33.5** | 0.386 | **21.5** | 0.4841 | 1.5 | 352.666 | **10.9** | 364.798 | 7.8 | 380.765 | 3.8 | 247.667 | -6.0 | 308.111 | -31.9 | 351.056 | -50.3 |
| **20** | DP01312SC | 0.29055 | **40.9** | 0.3713 | **24.5** | 0.43248 | 12.0 | 340.645 | **13.9** | 362.651 | 8.4 | 375.951 | 5.0 | 188.667 | 19.2 | 251.778 | -7.8 | 303.778 | -30.0 |
| **21** | HTS03456SC | 0.27456 | **44.2** | 0.4391 | 10.7 | 0.42156 | 14.3 | 348.891 | **11.8** | 380.403 | 3.9 | 380.881 | 3.7 | 190.222 | 18.6 | 263.833 | -12.9 | 254.278 | -8.8 |
| **22** | **HTS06879SC** | 0.489 | **30.6** | 0.57985 | 17.7 | 0.64277 | 8.7 | 340 | **10.9** | 362 | 5.4 | 370 | 3.0 | 339 | 10.0 | 353 | 6.4 | 401 | -6.5 |
| **23** | **HTS07222SC** | 0.46515 | **33.9** | 0.59259 | 15.9 | 0.61829 | 12.2 | 344 | 9.9 | 354 | 7.3 | 365 | 4.6 | 307 | 18.5 | 359 | 4.8 | 419 | -11.4 |
| **24** | **JP00899SC** | 0.49023 | **31.9** | 0.57998 | 19.5 | 0.67144 | 6.8 | 350 | **11.3** | 365 | 7.6 | 394 | 0.3 | 353 | 15.5 | 393 | 5.9 | 417 | 0.1 |
| **25** | **SPB06618SC** | 0.48773 | **30.1** | 0.59504 | 14.7 | 0.56582 | 18.9 | 370 | 5.5 | 367 | 6.2 | 357 | 8.8 | 314 | 25.0 | 390 | 7.0 | 391 | 6.7 |

| | | Puncta/NLS | | | | | | Vesicle Average Intensity | | | | | | mCherry NLS | | | | | |
|---|---|---|---|---|---|---|---|---|---|---|---|---|---|---|---|---|---|---|---|
| | | 10uM | | 3uM | | 1uM | | 10uM | | 3uM | | 1uM | | 10uM | | 3uM | | 1uM | |
| Hit No | **Enamine CAT_No** | AVE | % | AVE | % | AVE | % | AVE | % | AVE | % | AVE | % | AVE | % | AVE | % | AVE | % |
| **26** | **Z198022656** | 0.32155 | **48.7** | 0.45333 | **27.6** | 0.49726 | **20.6** | 327 | 8.7 | 337 | 6.0 | 343 | 4.5 | 290 | 14.8 | 342 | -0.5 | 350 | -2.8 |
| **27** | **Z223817830** | 0.41479 | **40.0** | 0.63704 | 7.8 | 0.62615 | 9.4 | 350.906 | 7.2 | 366.085 | 3.2 | 370.744 | 2.0 | 558.5 | 7.5 | 638.333 | -5.7 | 658.833 | -9.1 |
| **28** | **Z31189063** | 0.36566 | **47.1** | 0.51748 | **25.1** | 0.61892 | 10.4 | 336.793 | **11.0** | 354.127 | 6.4 | 367.715 | 2.8 | 639.889 | -5.9 | 640.333 | -6.0 | 665.833 | -10.2 |
| **29** | **Z228869724** | 0.31816 | **44.8** | 0.51922 | 9.9 | 0.56497 | 2.0 | 338.045 | **9.2** | 360.334 | 3.2 | 368.063 | 1.1 | 339.222 | 19.3 | 448.444 | -6.7 | 438.389 | -4.3 |
| **30** | **Z387344412** | 0.09142 | **84.1** | 0.24639 | **57.3** | 0.45115 | **21.7** | 313.549 | **15.7** | 329.703 | **11.4** | 354.445 | 4.8 | 323.5 | 23.0 | 435.389 | -3.6 | 443.333 | -5.5 |
| **31** | **Z102923060** | 0.42117 | **33.8** | 0.58053 | 8.8 | 0.59457 | 6.6 | 338.483 | **10.6** | 359.966 | 5.0 | 368.484 | 2.7 | 384.389 | 22.0 | 500.445 | -1.5 | 539 | -9.4 |
| **32** | **Z53051092** | 0.34124 | **46.4** | 0.49572 | **22.1** | 0.59518 | 6.5 | 329.238 | **13.1** | 344.445 | **9.1** | 360.453 | 4.8 | 411.278 | 16.6 | 479.667 | 2.7 | 523.555 | -6.2 |
| **33** | **Z203923506** | 0.39583 | **37.4** | 0.53333 | 15.7 | 0.55682 | 12.0 | 332.865 | **12.1** | 349.707 | 7.7 | 359.354 | 5.1 | 349.944 | 20.5 | 431.611 | 1.9 | 438.611 | 0.3 |

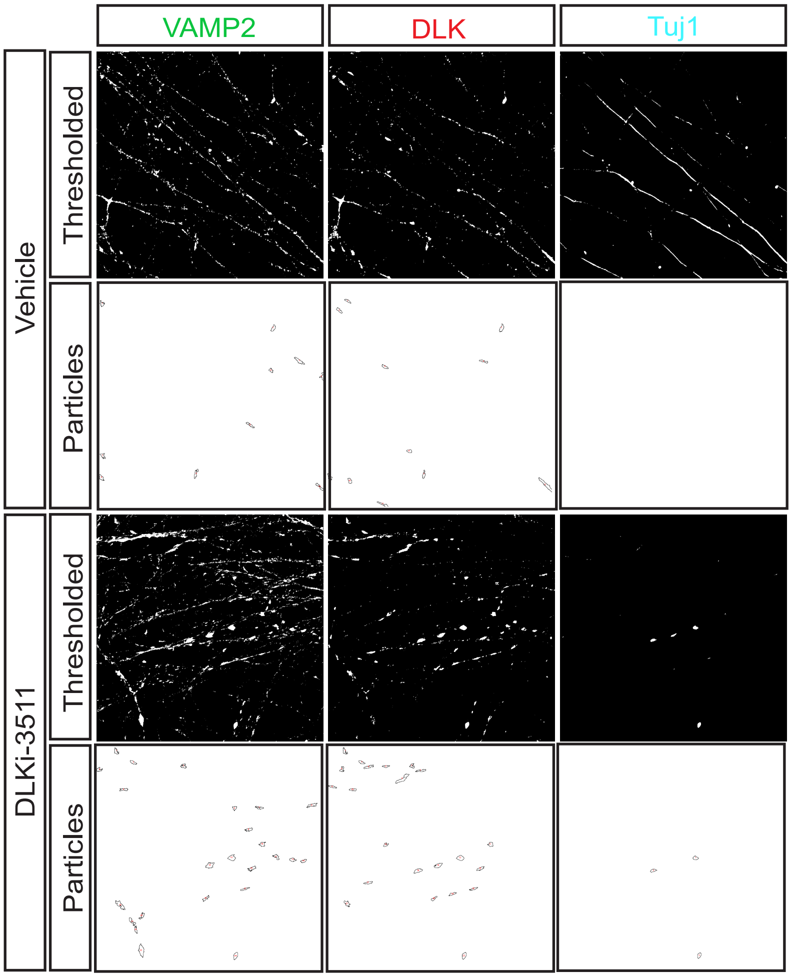


**Figure S1: Example of thresholding and particle count analysis images related to Figure 1.** Images of individual channels from Figure 1B were subjected to auto-thresholding and particle count analysis, as described in Methods. Images of these image processing steps, including outlined puncta/accumulations (‘Particles’), whose area is then quantified relative to the total thresholded area for each given signal, are shown

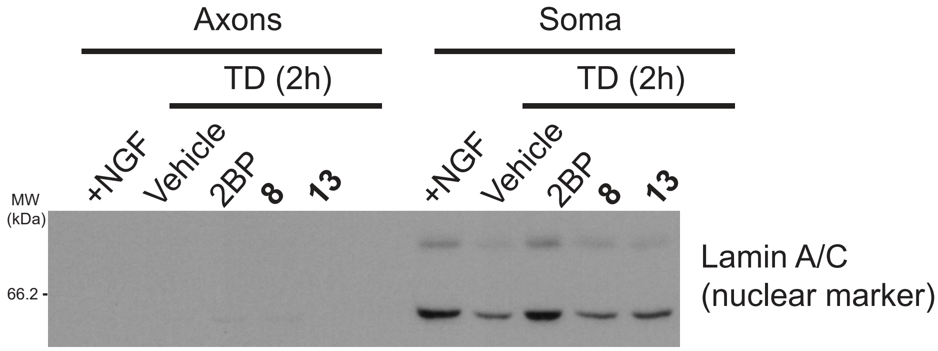


**Figure S2: Confirmation of fidelity of axonal preparation for Fig 2C.** Lysates of axonal fractions used in Fig 2C were subjected to SDS-PAGE and western blotting with Lamin A/C antibody (nuclear marker) side-by-side with somal fractions from the same cultures. Lamin A/C are essentially absent from axonal fractions.

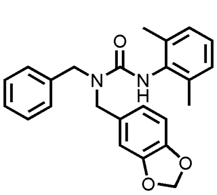

**1**

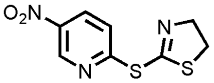

**2**

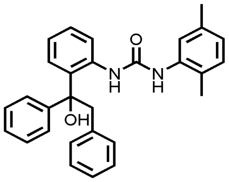

**3**

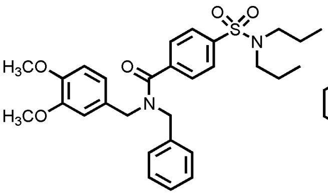

**4**

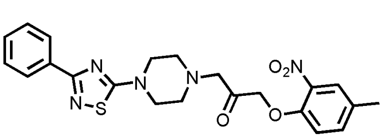

**5**

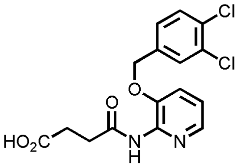

**6**

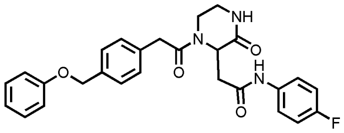

**7**

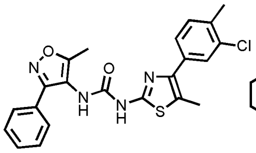

**8**

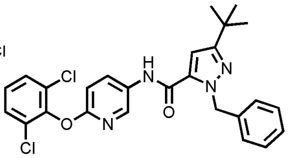

**9**

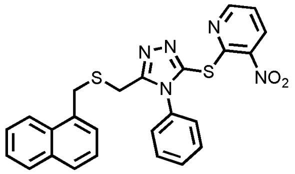

**10**

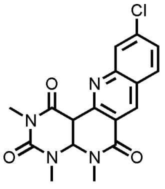

**11**

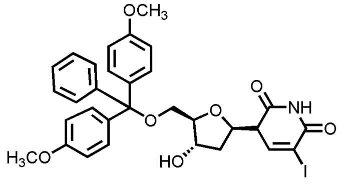

**12**

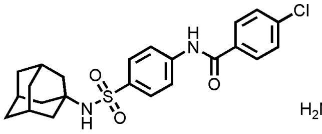

**13**

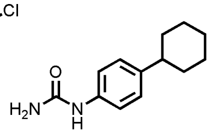

**14**

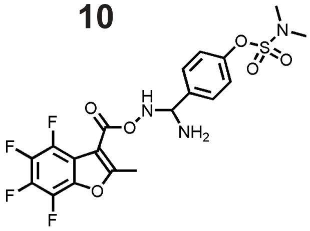

**15**

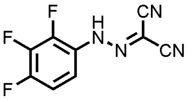

**16**

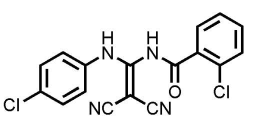

**17**

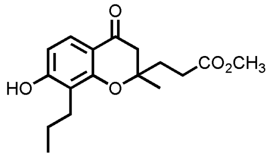

**18**

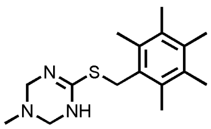

**19**

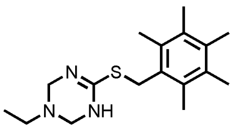

**20**

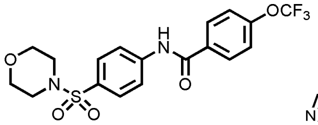

**21**

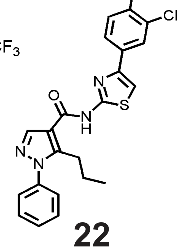

**22**

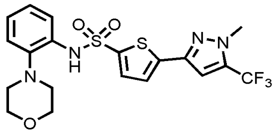

**23**

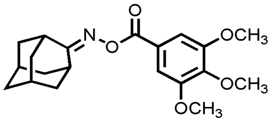

**24**

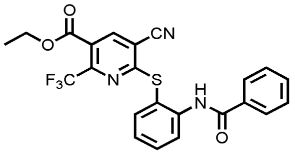

**25**

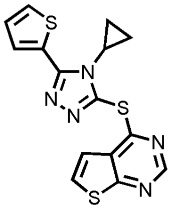

**26**

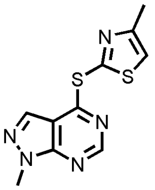

**27**

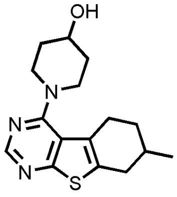

**28**

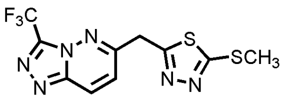

**29**

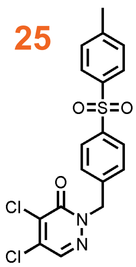

**30**

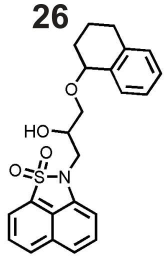

**31**

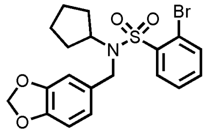

**32**

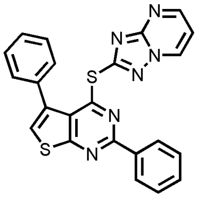

**33**

**Figure S3: Structures of compounds used in TD-induced c-Jun phosphorylation assay in Figure 4.** Identifying numbers for compounds followed up in neurodegeneration assay in Figure 5 are highlighted in orange.

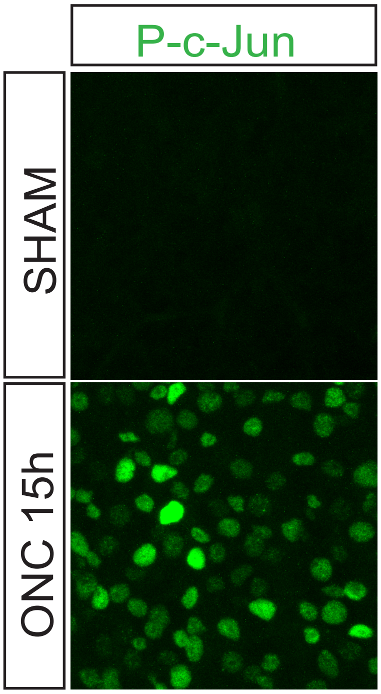


**Figure S4: Confirmation of acute, ONC-induced c-Jun phosphorylation.** Mice were subjected to ONC or sham injury and were fixed and perfused 15h later. Immunostaining of retinas with the indicated antibody confirms robust ONC-induced c-Jun phosphorylation (p-c-Jun) at this time point.
